# Decoupling the molecular regulation of perenniality and flowering in bulbous barley (*Hordeum bulbosum*)

**DOI:** 10.1101/2021.11.11.468190

**Authors:** Dana Fuerst, Bar Shermeister, Tali Mandel, Sariel Hübner

## Abstract

Global crop production is being challenged by rapid population growth, declining natural resources, and dramatic climatic turnovers. These challenges have prompted plant breeders to explore new ventures to enhance adaptation and sustainability in crops. One intriguing approach to make agriculture more sustainable is by turning annual systems into perennial which offers many economic and biodiversity-friendly benefits. Previous attempts to develop a perennial cereal crop employed a classical breeding approach and extended over a long period with limited success. Thus, elucidating the genetic basis of perenniality at the molecular level can accelerate the breeding process.

Here, we investigated the genetic basis of bulb formation in the barley congener species *Hordeum bulbosum* by elucidating the transcripts presence/absence variation compared with other annual species in the *Poaceae*, and a differential expression analysis of meristem tissues. The PAV analysis recaptured the expected phylogeny and indicated that *H. bulbosum* is enriched with developmental and disease responsive genes that are absent among annual species. Next, the abundance of transcripts was quantified and allowed to identify differentially expressed genes that are associated with bulb formation pathways in addition to major circadian clock genes that regulate flowering. A first model for the bulb formation pathway is suggested and include developmental and starch biosynthesis genes. To the best of our knowledge this is the first transcriptome developed for *H. bulbosum* and the first attempt to describe the regulation of bulb initiation in cereals at the molecular level.

## INTRODUCTION

It is now recognized that agriculture can be made far more sustainable by transitioning many annual agricultural systems to perennial (Ryan et al., 2018). Perennials are year-round crops harvested multiple times over several seasons, a feature that offers many environmental and economic benefits. Unlike annual crops, perennials are thriftier and improve soil structure and water retention capacity, contribute to increased mitigation practices to cope with climate change, and promote biodiversity and ecosystem functions (Kantar et al., 2016; Lundgren et al., 2020). Among annual crops that are cultivated across over 70% of global croplands, many staple crops could potentially be transferred to a perennial life habit by hybridization and other genomic engineering techniques (Hübner and Kantar 2021). Generally, there are two main approaches to develop a new perennial crop: *de novo* domestication of a wild perennial plant, and introgression of perennial traits into an annual crop through hybridization practices (Cui et al., 2018; Kantar et al., 2018). Several attempts to develop a perennial cereal crop were made over the years with little commercial success. For example, efforts to develop perennial wheat varieties through hybridization with wild relatives were successful in generating stable amphiploids (Amstrong, 1936; Larkin et al., 2014), yet grain yield declined quickly under field conditions. Thus, introgression of perenniality into an annual crop is a challenging long endeavor. Targeting crosses and directing selection based on a deep understanding of the molecular regulation of the perenniality traits can potentially accelerate this tedious process (Kantar et al., 2018). Recent developments in biotechnology have prompt new methods to introduce a trait of interest into a crop through genome editing. This technology can be used for *de novo* domestication of wild species by targeting genes that regulate domestication syndrome traits including seed shattering and awn length (Yu et al., 2021), fruit size and number, and the nutritional value (Gasparini et al., 2021). Another approach, considers perenniality as a syndrome which includes a variety of interacting traits, thus understanding the underlying molecular mechanism is necessary to allow breeding efforts to focus on specific components of perenniality (Lundgren and Marais 2020).

Barley (*Hordeum vulgare*) ranks fourth among cereal crops in cultivated area (www.fao.org/faostat) and is known for its enhanced adaptation to a wide range of environments (Haberer et al., 2015). Very few attempts were made so far to breed for perenniality in barley although there are perennial species in the genus that are cross-compatible with cultivated barley.

*Hordeum bulbosum* is a wild cereal species that has diverged from barley 4 million years ago and is highly abundant in the Mediterranean region (Brassac et al., 2015). Despite the morphological similarity between barley and *H. bulbosum*, the latter is a perennial species merit to its bulb organ, which allows to pass the dry summer season by entering a dormancy period. In the late fall, when the days are getting shorter, temperature declines and the first rain of the season, the new *H. bulbosum* leaves sprout from regenerating buds that are located at the periphery of the bulb organ. After emergence, the *H. bulbosum* plant continues to grow and establish until the end of December when days are getting longer and signal the plant to shift from vegetative growth to flowering and to form a new bulb organ (Koller et al., 1960). Previous studies in anion (*Allium cepa L*.) indicated that the transition from short to long days induce bulb formation and development through the regulation of FT homologs and other flowering regulating genes (Lee et al., 2013). However, little is known about the regulation of bulb formation in *H. bulbosum* and the underlying molecular mechanism remains largely unknown. To address this, advanced genomic infrastructure and proper molecular tools adjusted for *H. bulbosum* are mandatory, yet very few tools were developed so far for this non-model species. Recently, a reference draft genome was developed for a haploid *H. bulbosum*, but this resource is highly fragmented and un-annotated, thus the representation of the gene space is partial (Wendler et al., 2017). Another efficient approach to identify genes of interest in a non-model organism is RNA sequencing which allows to efficiently develop a reference transcriptome and test for the expression of genes in response to a signal (Costa-Silva et al., 2009). Assembling *de novo* a transcriptome for *H. bulbosum* has many benefits and can significantly improve the gene space representation and the identification of candidate genes involved in a perennial organ formation and development.

Here, we describe the assembly of a reference transcriptome for *H. bulbosum* and the analysis of the presence/absence variation and differential gene expression to identify candidate genes that are involved in the regulation of bulb formation. We hypothesize that the development of a bulb organ in *H. bulbosum* is initiated in the shoot meristem during the vegetative stage in response to transition from short-day to long-day regime. We show that bulb formation is regulated by genes and pathways that are coupled with the circadian clock and flowering but the experimental design used allowed to partially deconvolute them. We further explore the set of candidate genes identified and suggest a molecular model for the regulation of bulb initiation in *H. bulbosum*.

## MATERIALS AND METHOD

### Plant material

A controlled experiment was set from a random batch of seeds sampled at Alon HaGalil in northern Israel (32°45’35.2”N 35°13’30.5”E), from single spikes of *H. bulbosum* and *H. spontaneum* plants. Selected seeds were sown in planting trays, and placed in a cold (4°C) dark room for 15 days to break dormancy. Germinated seedlings were transferred into a growth-room under short-day regime (SD; 8h:16h light:dark), and constant temperature of 16°C. After an establishment period of two weeks, seedlings were transferred into 3-liter pots containing a soil mixture (“GREEN 90”, Ben-Ari Ltd., Israel) and slow release fertilizer (Osmocote, Everris International B.V. Heerlen, The Netherlands). After nine weeks in growth-room under SD regime, plants of both species were divided randomly between three groups in accordance with the three treatments in the experiment (T_0_, T_2_, T_4_). Each treatment was set in three biological replicates, thus a total of 9 plants of each species (total 18 plants) were included in the experiment. Shoot meristem tissues were sampled from the first group (T_0_) before transition to long-day (LD) regime and were immediately frozen in liquid nitrogen. After sampling the T_0_ plants, the conditions in the growth-room were shifted to LD regime (16h:8h, light:dark) and shoot meristems were sampled from the T_2_ plants after 24 hours. After additional 96 hours in LD, shoot merisms were sampled from the third group of T_4_ plants.

For the reference transcriptome assembly, three *H. bulbosum* plants were grown under SD for three weeks and then transferred to LD regime until flowering and development of a mature bulb organ. A total of six tissues were sampled for RNA extraction and sequencing including bulb, anthers, stigmas, leaf, embryo and shoot meristem. For embryo extraction, a single seed was soaked in water for two hours to soften the seed and allow easy removal of the embryo. Altogether, twenty-four tissue samples were obtained and immediately frozen in liquid nitrogen and stored in −80°C until extraction of RNA.

### RNA extraction, library preparation and sequencing

Total RNA was extracted from each of the 24 samples (9 *H. spontaneum* and 15 *H. bulbosum*) using the RNeasy Plant Mini Kit (QIAGEN cat No./ID 74904) following the manufacturer protocol. For embryo RNA extraction, we used 500 µl RLT buffer and 10µl β-ME, and for the leaf tissue 500 µl RLT buffer and 5µl β-ME as recommended by the manufacturer. To confirm the integrity and quality, total RNA was inspected in both NanoDrop 1000 spectrophotometer and agarose gel electrophoresis. An OD 260/280 ratio of 2.0 and RNA concentration of 100 ng/µl were set as a minimum threshold for adequate quality and quantity. Sequencing libraries were constructed and sequenced at the Technion Genome Center (Haifa, Israel). RNA-Seq libraries were prepared using the TruSeq RNA Library Prep Kit v2 (Illumina Inc., USA) following manufacturer instructions and sequenced on six lanes of HiSeq-2500 Illumina machine. To avoid a lane bias in the expression profiling analysis, samples were pooled and the sixth portion of the pool was sequenced on each of the six lanes. The quality of the sequenced reads was inspected with the software FastQC v.0.11.5 (Andrews, 2010) and low quality reads were trimmed and adapters were removed using the default parameters in Trimmomatic v.0.32 (Bolger et al., 2014). Finally, cleaned and high-quality reads were re-inspected using FastQC.

### Transcriptome assembly and annotation

To develop a comprehensive and representative transcriptome for *H. bulbosum*, 15 RNA-Seq libraries obtained from roots, bulb, leaf, floral reproductive tissues (anthers and stigma) and shoot meristems were sequenced on a HiSeq-2500 Illumina machine. The transcriptome was assembled *de novo* with the Trinity assembler v1.8 (Grabherr et al., 2011) using the reads generated from all *H. bulbosum* tissues. Briefly, Trinity performs a full transcriptome assembly process in three main stages and include partitioning the data, clustering and graphs construction, and tracing paths in the constructed graphs in parallel. At the end of this process, a linear sequence is obtained for each transcript isoform.

To annotate the transcriptome we used the Trinotate pipeline v-3.1.1 (Bryant et al., 2017). Prediction of open reading frame (ORF) in the assembled genes was conducted with TransDecoder v-5.5.0 (http://transdecoder.github.io). Homolog proteins detection was conducted with blastx and blastp against the SwissProt and UniProt (Boeckmann et al., 2005; the UniPort Consortium, 2021) databases and an e-value cut-off of 10^−5^. To identify conserved protein domains, trans membranal regions and rRNA genes, the HMMER toolkit v-3.1b2 (Eddy, 1998) was used against PFam database (Finn et al., 2014). Signal peptides were predicted using SignalP (Petersen et al., 2011). To filter potential contamination in the transcriptome, transcripts that were annotated to species outside the *Spermatophyta* super-division (seeded plants) were excluded from the dataset. To further evaluate the quality of the assembly, cleaned reads from each of the sequenced libraries were aligned back to the filtered transcriptome using bowtie2 v2.3.5.1 (Langmead et al., 2009) and the alignment statistics was calculated with samtools v1.9-92 (Li et al., 2009). To evaluate the completeness of the transcriptome, a BUSCO analysis was preformed (Simão et al., 2015) using the *Poales* database and the transcriptome mode. Only longest isoform per gene was inspected to minimize the false identification of duplicated genes.

Finally, the obtained transcriptome was compared with the genomes of *Hordeum spontaneum* (Jayakodi et al., 2020), *Trititcum aestivum* (Appels et al., 2018), *Oryza sativa* (Wang et al., 2018), and *Brachypodium distachyon* (Vogel et al., 2010) and presence absence variation among species was analyzed. We extracted the longest isoform from each gene to avoid the complex comparison between a transcript and the gene model as annotated in different species. The extracted isoform was then searched in the available draft genome of *H. bulbosum* to exclude genes that are specific to the individual that was sequenced and the assembly methodology in each platform. The genes presence/absence among species was determined using blastn where no-hit of a gene was considered as absent.

### Differential expression analysis

To identify candidate genes involved in the regulation of bulb formation in *H. bulbosum* a differential expression (DE) analysis was performed using the RNA-Seq data generated from shoot meristem tissues before and after transition from SD to LD photoperiod. Previous studies on bulb physiology and development (Ofir et al., 1974) have shown that the day-length signal is intensified 8-10 weeks after germination, thus plants were kept in SD regime for this period of time. The experiment included three treatment groups: before the transition to LD regime (T_0_), 24 hours after transition to LD regime (T_2_), and 96 hours after the transition to LD (T_4_). For each group, three biological replicates were sampled for RNA extraction and sequencing. Obtained data was trimmed and cleaned with Trimmomatic and only high-quality reads were aligned with Bowtie2 to the transcriptome. To quantify the expression profile of genes in each sample, alignment files were processed with RSEM v.1.2.31 (Li et al., 2011), and genes with differentially expressed profile between treatments were detected with the R package DESeq2 v1.22.2 (Love *et al*., 2014). To correct the high false discovery rate (FDR) due to multiple testing, we used the Benjamini-Hochberg correction with a cutoff of FDR ≤ 5%. No filtering threshold was set for the expression fold change (log_2_-FC) between treatments to avoid the exclusion of mild differences which is common for regulating genes such as transcription factors (Sha’ar-Moshe et al., 2015).

### Gene ontology analysis

Gene ontology (GO) analysis was conducted to identify enriched biological processes among the different tissues that were used to assemble the transcriptome, among genes showing presence/absence variation (PAV), and among genes that were differentially expressed in response to transition from SD and LD regimes. GO terms were extracted from the annotation file generated for the transcriptome and used for enrichment analysis with the package topGO v3.11 (Alexa et al., 2006) in R. The GO analysis was performed with default parameters and statistical significance was determined with Fisher exact test. The *p*-values reported were not adjusted following the recommendations of the package authors. Heatmaps were generated with ‘gplot’ v3.1.1 (Warnes et al., 2009) for the top significant GOs terms.

## RESULTS

### The *H. bulbosum* transcriptome

A total of 140 billion base-pairs from 15 tissue samples were processed in order to assemble the *H. bulbosum* transcriptome after removing 2-4% of reads per sample due to unsatisfying quality. The total length of the obtained transcriptome was 650 Mbp comprising of 707,824 transcripts (Table S1). Following the assembly, transcripts were annotated and filtered to include only genes that are present in the *Spermatophyte* super-division database. This filtering approach allowed to avoid potential contamination while maintaining information from species outside the *Poacea* that may be relevant for the identification of bulb regulating genes (e.g. onion, garlic). A total of 138,696 transcripts remained after filtering, of which 42,045 are unigenes with a contig N50 of 2,758 bp. Reads from all libraries of *H. bulbosum* were aligned to the filtered transcriptome and the average alignment rate was 59% (Table S2, Figure S1).

To evaluate the completeness of the assembled transcriptome, BUSCO analysis was performed using the *Poales* genes database (4,896 BUSCOs) and the transcriptome mode. Altogether, 3,117 BUSCO genes were detected of which 92% are complete genes and 84% are single copy genes. Next, the assembled transcriptome was compared with the available reference draft genome for *H. bulbosum* using the BUSCO pipeline and the *Poales* genes database. Expectedly, more genes were detected in the draft genome (4,119) than in the transcriptome because the later was targeted to specific tissues and timing. Interestingly, once unrepresented genes were excluded, the number of complete genes was higher (92%) and the number of duplicated genes was lower (87%) in the transcriptome compared with the draft genome (Figure 1, Table S3). These results emphasize the advantage of the transcriptome in representing targeted genes for expression profiling and analysis.

**Figure 1.**
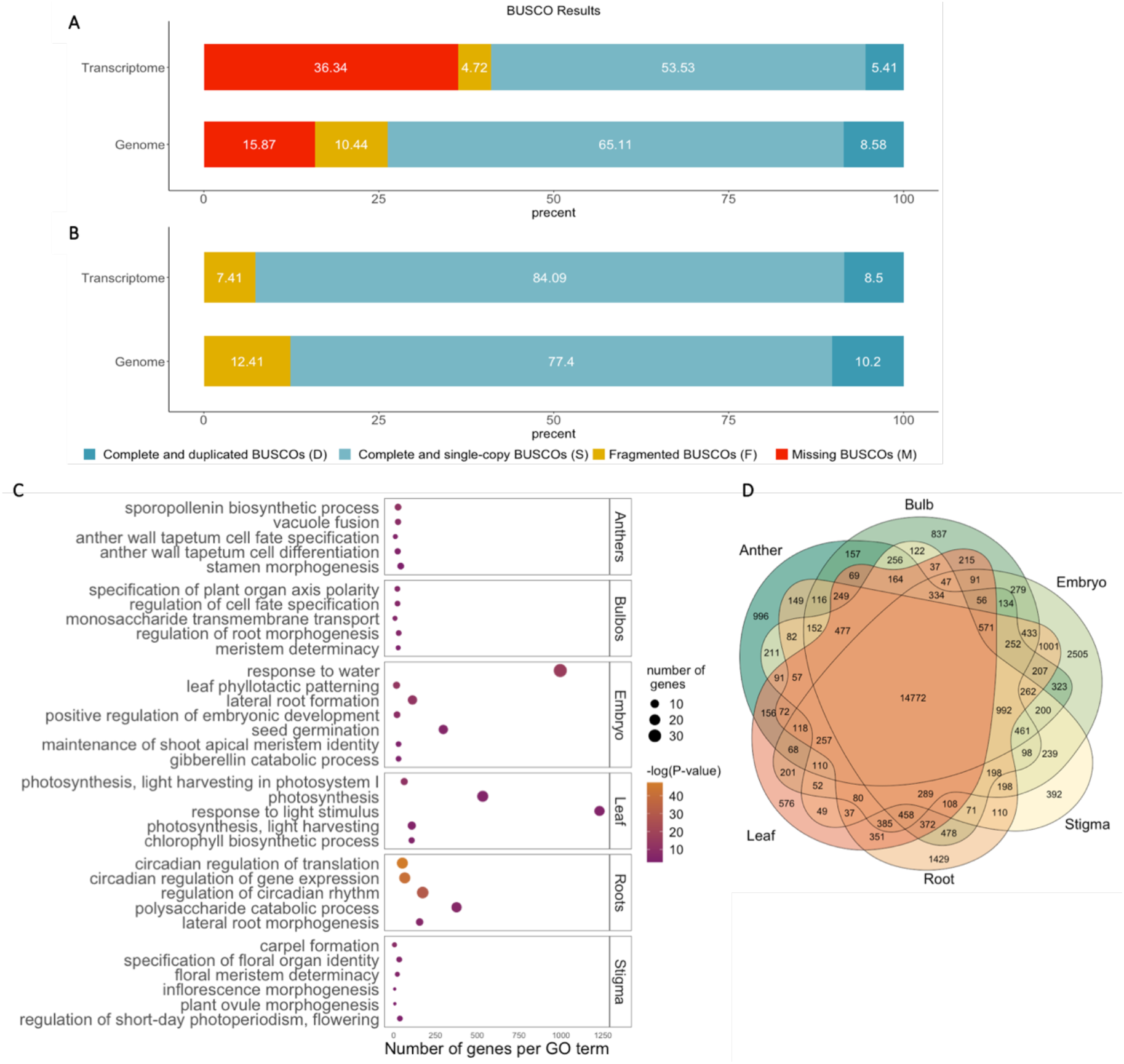
Evaluation and analysis of H. bulbosum transcriptome. **A**,**B)** Proportion of BUSCOs in the de novo transcriptome assembly compared with the draft genome of H. bulbosum including all BUSCO genes (A), and after removing missing BUSCOs (B). **C)** Significantly enriched GO terms in each tissue that was included in the transcriptome assembly. The circle size is proportional to the number of genes associated with in each GO term. **D**) Venn diagram of shared and unique genes expressed among tissues.

To further explore the transcriptome composition and the expression profile in each tissue (leaf, root, flower organs, bulb and embryo), a subset of 34,282 genes with a minimum coverage of 2X in at least one tissue were extracted. Genes that were expressed only in shoot-meristems were excluded from this analysis due to over-representation among sequenced libraries (9 libraries). The number of genes detected in each tissue was quantified and indicated that 43% of the expressed genes are shared among all 6 tissues, and 19% are tissue-specific (Figure 1). High rate of tissue-specific genes was observed among embryo (2,505), followed by roots (1,429), anthers (996), bulb (837), leaf (576), and stigma (392), after confirming that the number of genes detected in a tissue is not biased by the amount of RNA extracted (*r* = 0.29, *p* = 0.57; Figure S2). To further explore the underlying function of genes that were expressed in different tissues or tissue-specific genes, a gene ontology (GO) enrichment analysis was conducted (Table S4). Among GO categories, 19 were significantly enriched across all tissues and included response to stimulus, developmental process and biological regulation. Within tissues, GO categories that were significantly enriched in a specific tissue included biological processes that are identified with the corresponding tissue (Figure 1C). For example, leaf tissue was enriched with photosynthesis processes including light harvest in photosystem-I (GO:0009768, *p* = 1.70×10^−5^) and chlorophyll biosynthesis (GO:0015995, *p* = 8.17×10^−3^), embryo tissue was enriched with germination processes (GO:0009415, *p* = 9.40×10^−8^), and phyllotaxis (GO:0060772, *p* = 2.30×10^−6^). A list of GO terms identified in each tissue is provided in the supplementary materials (Table S4).

The ability to generate a perennial organ among *Triticeae* species is rare, thus gene gain or loss dynamics may underlie the process of bulb formation and development in *H. bulbosum*. To test this and identify genes that are present in *H. bulbosum* and absent among annual grass species, the presence/absence variation (PAV) between *H. bulbosum, H. spontaneum, T. aestivum, B. distachyon and O. sativa* was investigated. Among the 42,045 assembled genes in the transcriptome, 2,591 were absent from the draft genome and were excluded from the PAV analysis. All remaining transcripts were compared to all other species to identify and quantify the level of homology for each transcript using blastn. A gene was considered absent in a species only if no-hit was obtained. This conservative approach allows to target, with high confidence, genes in *H. bulbosum* that are absent in the other annual species albeit with some level of false negative (undetected genes). A phylogenetic tree constructed from the PAV across all genes supported the expected topology among the *Triticeae*. The lowest rate of genes shared with *H. bulbosum* was observed for *O. sativa* (47% of *H. bulbosum* genes) which has diverged from *H. bulbosum* 50 MYA, and highest rate (91%) was observed for *H. spontaneum* which has diverged from *H. bulbosum* 4 MYA (Figure 2A,C). Among the 39,454 genes for which PAV was detected, 154 genes were unique to *H. bulbosum* and 626 were present only within the *Hordeum* genus. Among unique genes to *H. bulbosum*, number of disease resistance genes were detected including RGA-like genes which are involved in plant-pathogen interaction. Among the genes detected in the transcriptome assembly and absent from all genome assemblies (including the *H. bulbosum* draft genome), several organ development genes were detected including the 14-3-3-like gene which promote tuberization in potato (Teo et al., 2017). Interestingly, 14-3-3-like was also significantly differentially expressed in response to the transition to LD regime in *H. bulbosum* but was absent from the annual *H. spontaneum*.

**Figure 2.**
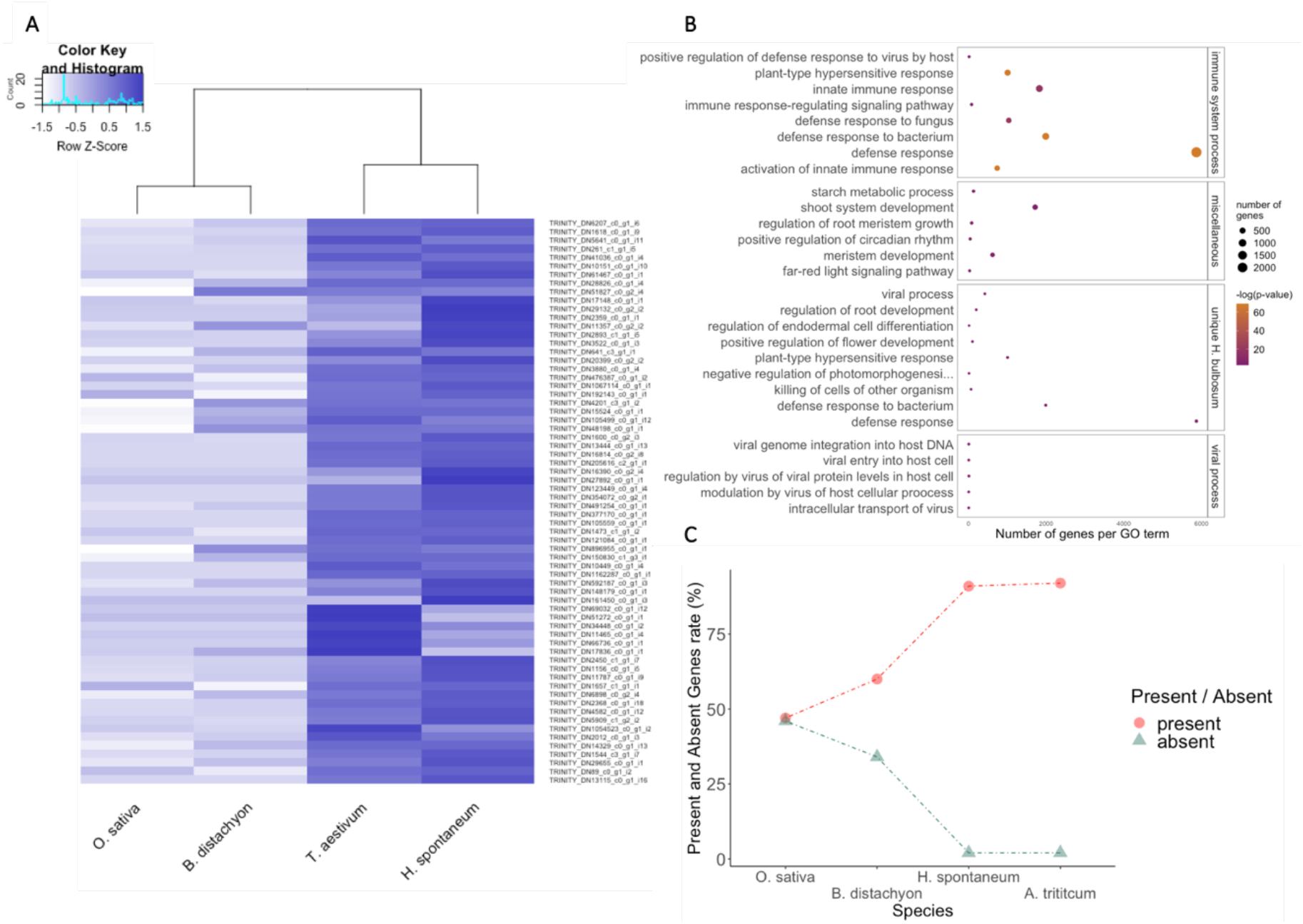
Presence/absence variation (PAV) analysis for H. bulbosum and annual species of the Poaceae. **A)** Heatmap of PAV genes in H. bulbosum compared with O. sativa, B. distachyon, H. spontaneum, and T. aestivum using a normalized bit-score values obtained from blastn analysis. Genes with no-hit were given a score of 0 and darker colors indicate higher homology. **B)** Significantly enriched GO terms among genes that were identified in H. bulbosum but are absent in at least one species or unique to H. bulbosum. **C)** Percent of P/A genes among different species.

To further investigate the biological process that underlie the detected PAV genes, a GO enrichment analysis was conducted. A total of 111 GO terms were significantly enriched among genes that were detected only in *H. bulbosum* and absent in annual species and included response to biotic stress (GO:0050832, *p* = 1×10^−30^; GO: 0046718, *p* = 9.9×10^−3^), sugar accumulation, starch metabolism and organ development (GO:0005982, *p* = 2.8×10^−2^; GO: 0042753, *p* = 7.9×10^−5^; GO: 0010082, *p* = 1.4×10^−3^; GO: 0048367, *p* = 3×10^−2^) (Figure 2B, Table S5).

### Identification of candidate genes involved in bulb formation

Previous work suggested that the bulb initiation signal is intensified at the meristem after 8-10 weeks in short-day regime (Ofir et al., 1974). Therefore, meristem tissues were sampled 10 weeks after germination (T_0_) and then were transitioned to LD regime. Following the transition, meristem tissues were sampled after 24 (T_2_) and 96 (T_4_) hours under LD and RNA was extracted and sequenced (Figure 3A). An average of 92 million paired-end reads were obtained for each sample and were aligned to the assembled transcriptome yielding an average of 60% and 30% properly aligned and uniquely mapped reads, respectively (Table S2). The high rate of multi-mapped reads with low mapping quality was further investigated using a subset of 1000 reads that were randomly sampled and searched against the NCBI non-redundant database. The analysis indicated that the high rate of multi-mapped reads is attributed to the large number of duplicated genes in the assembled transcriptome. Finally, a total of 36,929 genes were detected among the meristem tissues with an average coverage of 66% and an average depth of 38X (Table S6).

**Figure 3.**
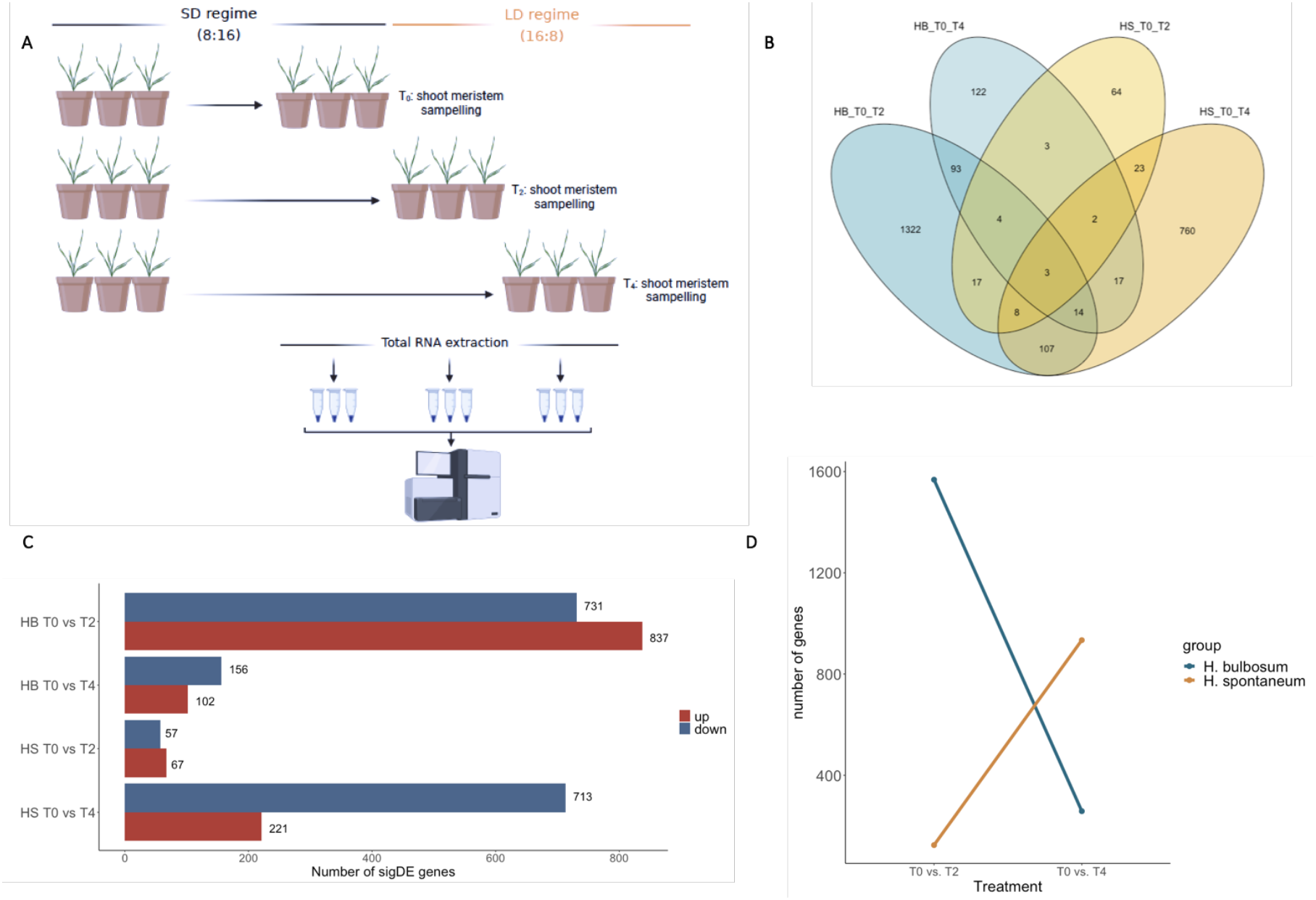
Differential expression analysis of meristem tissues in response to shift in day-length regime. **A**) A scheme diagram of the experiment. **B**) A Venn diagram for DEGs detected in H. bulbosum and H. spontaneum. **C)** Counts of up and down DEGs in H. bulbosum (HB) and H. spontaenum (HS). **D)** Total number of DEGs in H. bulbosum and H. spontaneum along time/treatment.

Next, the expression profile at different treatments and timepoints was investigated. A PCA conducted for the different samples using the expression profile of all genes confirmed the assignment of samples to treatments (Figure S3). A major challenge in the experiment was that LD signal initiates the formation of a bulb organ, but as in other species, also the transition to flowering. To decouple these two processes and target genes that underlie the regulation of bulb formation, the experiment was conducted also for the annual species *H. spontaneum* following the same experimental setup. We hypothesize that unlike flowering regulating genes that are expected to be detected in both species, bulb formation and development genes will be detected only in the perennial species (*H. bulbosum*).

Following the transition of *H. bulbosum* from SD to LD (T_0_/T_2_), a total of 1568 significantly differentially expressed genes (DEGs) were detected of which 837 were upregulated and 731 were downregulated (Figure 3). After four days in LD (T_0_/T_4_) the effect has strongly declined and only 102 genes were upregulated and 156 were downregulated. In contrast to the expression profile observed in *H. bulbosum*, the transition from SD to LD regime in *H. spontaneum* had a mild effect after 24 hours and only 67 upregulated and 57 downregulated DEGs were detected. However, after four days in LD the effect was intensified and 221 upregulated and 713 downregulated DEGs were detected. These results indicate that the day length signal is indeed intensified in *H. bulbosum* due to the long period in SD (10 weeks) while in *H. spontaneum* the effect accumulates gradually after the transition to LD. Interestingly, flowering related genes were mainly detected in *H. bulbosum* after four days (T_4_) in LD similarly to the response observed in *H. spontaneum*, implying that bulb initiation genes responded to the transition to LD regime before flowering regulating genes (Figure 3E).

To further study the functional context of the detected DEGs in *H. bulbosum*, a GO enrichment analysis was conducted. The number of significantly enriched GO terms was consistent with the declining trend observed in the expression analysis, thus 183 and 46 enriched GO terms were detected for T_0_/T_2_ and T_0_/T_4_ respectively. Among the enriched GO terms in the T_0_/T_2_ and the T_0_/T_4_ comparisons, 25 terms are associated with flowering (GO:0045595, *p* = 0.0038; GO:0009908, *p* = 0.0036), response to photoperiod (GO:0048577, *p* = 0.0186), and bulb formation (GO:0005986, *p* = 0.0157).

To further explore the genetic basis of bulb initiation and development, a list of DEGs that are associated with significantly enriched GO terms was extracted (Table S7,S8). Generally, three phases are expected in the process of bulb formation: response to the LD signal, morphogenesis and organ development, and accumulation of sugars in the bulb organ. Among DEGs that underlie enriched GO terms, we detected CONSTANS (logFC = 1.48796, FDR = 4×10^−4^), GIGANTEA (logFC = −1.61, FDR = 1.49×10^−7^), and the pseudo-response regulator (PRR5) gene (logFC = −3.09, FDR = 1.71×10^−17^) which interact in response to day-length and signal transduction. In addition, we detected the morphogenesis and development regulating genes constitutive photomorphogenesis 9 (COP9) (logFC = 0.6096, FDR = 0.001) and MADS-box16 (logFC = 4.33, FDR = 2×10^−4^), and the fructose accumulation regulating gene SWEET17 (logFC = 2.11, FDR = 0.005).

In *H. spontaneum*, the response to LD was more prominent 96 hours after the transition to LD with 65 and 123 significantly enriched GO terms in the T_0_/T_2_ and T_0_/T_4_ respectively. Among the GO terms detected in the T_0_/T_4_, 13 are associated with the response to the LD signal and initiation of flowering (Table S9). GO terms that comprise DEGs that were also detected in *H. bulbosum* included MADS-box18, CONSTANS and PRR95, thus these genes are likely associated with initiation of flowering rather bulb formation and development.

## DISCUSSION

Despite the many benefits of modern agriculture, the associated environmental impact must be reduced to meet current global efforts to mitigate climate change. Thus, agriculture must adopt a more sustainable strategy to face the rising demand for food while natural rescues are declining. One way to reduce the negative environmental impact of agriculture is to shift crop plants from annual system to a perennial (Kantar et al., 2016, 2018; Lundgren and Des Marais, 2020; Mora et al., 2018). Compared with annual species, perennials tend to be more stress-tolerant and require less irrigation and environmentally harmful inputs including pesticides and fertilizers (Glover et al., 2010). Nevertheless, previous attempts to develop a perennial crop through classical breeding methods has extended over long periods due to challenging crosses between species, a sever genetic drag, and declining yield under field conditions (Kantar *et al*., 2016; Lundgren and Des Marais, 2020). Therefore, breeding towards perennial crops should become more targeted and incorporate biotechnological practices in the breeding process (Hübner and Kantar, 2021).

To allow an efficient targeted breeding, perenniality has to be accurately defined. From a broad perspective, perenniality can be considered as a syndrome that is comprised of many interacting traits including growth rate, carbon fixation, root system development, source-sink dynamics (Lundgren and Des Marais, 2020), and involve different plant organs. In this study, we defined perennially as the ability to form a bulb organ in *H. bulbosum* and followed two approaches to explore the underlying molecular mechanism.

### Identifying perennial life habit genes

The annual/perennial life habit has shifted in both directions along the evolution of many plant species (Heidel et al., 2016; Lundgren et al., 2020). This shift can be facilitated by a number of genetic mechanisms including gain or loss of “perenniality genes”, changes in the regulation of gene expression, or critical change in protein structure (Heidel et al., 2016). Along the evolutionary history of the *Hordeum* genus, perennial and annual life habits have shifted number of times independently (Blattner, 2009), and the course of life habit evolution among *H. bulbosum* and *H. spontaneum* remains elusive. Thus, perenniality may have evolved independently in *H. bulbosum* through a process of gene gain or loss. To test this, a presence/absence variation (PAV) analysis was conducted between *H. bulbosum* and other annual cereal species and indicated that many disease responsive and development genes evolved uniquely in *H. bulbosum* (Figure 2B). Interestingly, the gene 14-3-3-like was detected only in *H. bulbosum* and was previously found to promote storage organ formation in potato (Teo *et al*., 2017; Hannapel *et al*., 2017). This was further supported in the differential expression analysis, where 14-3-3-like was differentially expressed in response to the transition to LD regime in *H. bulbosum* but not in *H. spontaneum*.

### Decoupling the genetic mechanism involved in flowering and bulb formation

Another potential route for a perennial life habit to evolve is through modifications in genes expression that promote bulb formation and development. In *H. bulbosum*, bulb formation is induced by a shift in day-length similarly to onion and potato (Lee et al., 2013; Zhang et al., 2020), and develops from the bottom of the shoot meristem (Leshem, 1971). Unlike onion and potato where transition to flowering inhibits the development of the storage organ, in *H. bulbosum* flowering and bulb formation are coupled and occur simultaneously. To deconvolute bulb formation and flowering, a differential expression (DE) analysis was conducted using meristem tissues extracted before and after transition to long-day (LD) regime. The experiment was conducted in *H. bulbosum* and was compared with *H. spontaneum* where bulb initiating genes were not expected to be detected.

Comparison between the genes identified in *H. bulbosum* and known regulation pathways of flowering and storage organ development in potato, onion and Arabidopsis indicated that the mechanism is largely conserved. Generally, the initiation of flowering and bulb formation begins with reception of light signal in the leaf tissue and transmitted to the shoot meristem through flowering locus T (Hannapel et al., 2017; Lee et al., 2013). Here, the experiment was conducted only with shoot meristems, thus FT was not detected in *H. bulbosum* nor in *H. spontaneum*. Among the candidate genes identified in *H. bulbosum*, CO was upregulated and GIGANTEA was downregulated (Table S8). This expression profile promotes flowering in Arabidopsis (Putterill et al., 1995) but suppresses tuberization in potato (Abelenda et al., 2016, Hannapel et al., 2017). Moreover, the morphogenesis regulating gene COP9 which induces organ development (Chamovitz et al., 1996; Wei et al., 2003) was upregulated and so were several sugar transporters and starch biosynthesis genes including SWEET17 (Chardon et al., 2013).

This work sets the first milestone towards understanding the molecular mechanism of perenniality in *H. bulbosum*. Therefore, we cautiously suggest a general scaffold for the molecular mechanism that underlies the development of a bulb organ (Figure S4). The activity of COP9 triggers a rapid cells division followed by sugar biosynthesis and accumulation of starch (MADS-box, Cell division control protein 48, SWEET17 and Phosphoglycerate mutase-like protein 4). Most of the pathway seem to be conserved among *H. bulbosum* and *H. spontaneum*, thus breeding towards a perennial crop may be more achievable than expected.

## Conclusions

In this study, we assembled a reference transcriptome for *H. bulbosum* as a complementary infrastructure for the available draft genome. This platform has several advantages especially for targeting specific genes that are involved in bulb initiation and development. The PAV analysis of assembled transcripts and the differential expression analysis allowed to established the first milestone towards elucidating the molecular mechanism of bulb initiation and perenniality as was defined in *H. bulbosum*. The detected candidate genes should be further explored using genetic engineering techniques and later could potentially be applied in breeding programs to develop new perennial crop plants.

## Acknowledgements

This work was supported by the German-Israeli Foundation (GIF), Grant: No.I-2501-204.12-2018

## Notes

### Competing Interest Statement

The authors have declared no competing interest.

